# Adenosine modulates extracellular glutamate levels via adenosine A2A receptors in the delayed-ethanol induced headache

**DOI:** 10.1101/2020.10.02.324517

**Authors:** Nathan T. Fried, Christina R. Maxwell, Jan B. Hoek, Melanie B. Elliott, Michael L. Oshinsky

## Abstract

Identifying the mechanism behind delayed ethanol-induced headache (DEIH), otherwise known as the hangover headache, may provide insight into the mechanisms behind common headache triggers. Acetate was previously shown to be the key ethanol metabolite behind DEIH in the recurrent inflammatory stimulation (IS) rat model of headache. The reversal of trigeminal sensitivity following ethanol exposure with caffeine previously suggested a role of adenosine in DEIH. To characterize this, behavioral analysis and measurement of brainstem adenosine and glutamate with microdialysis and HPLC was performed while pharmacologically manipulating adenosine signaling in the IS and Spontaneous Trigeminal Allodynia (STA) rat models of headache. Blocking adenosine A_2A_ receptor activation with istradefylline or acetate transport into astrocytes with the monocarboxylate transporter competitive inhibitor, alpha-cyano-4-hydroxycinnamate (4-CIN), prevented acetate-induced trigeminal sensitivity. Blocking adenosine A_1_, A_2B_, and A_3_ receptor signaling did not prevent trigeminal sensitivity. Compared to control rats, IS rats had greater increases in extracellular adenosine and glutamate within the trigeminal nucleus caudalis (TNC) of the brainstem during local acetate perfusion. Blocking transport of acetate into astrocytes with 4-CIN prevented the increase in adenosine and glutamate. Blocking A_2A_ receptor activation prevented the increase in extracellular glutamate, but not adenosine in the TNC. These data are the first to demonstrate the physiological consequence of acetate on adenosinergic systems within trigeminal pain by suggesting that acetate-induced trigeminal sensitivity in DEIH is mediated by adenosine A_2A_ receptor activation which modulates extracellular glutamate levels in the TNC.

**Significance Statement:** It is unknown how several common headache triggers induce headache pain. Since migraineurs are more sensitive to these triggers, studying the mechanisms behind their effects may reveal unique migraine pathophysiology. In this study, we explored the common headache trigger, ethanol, which migraineurs are particularly sensitive to. When ethanol is ingested, its quickly metabolized to acetaldehyde and subsequently into acetate. We find that acetate increases brainstem adenosine and causes trigeminal sensitivity, which is exacerbated in the rat headache model. Blocking either acetate uptake or adenosine signaling prevents trigeminal sensitivity and brainstem glutamatergic signaling, suggesting that adenosine is involved in the hangover headache and that differences in acetate metabolism may account for the increased sensitivity to ethanol in migraineurs.

## INTRODUCTION

Alcohol’s capacity to induce the delayed ethanol-induced headache (DEIH), or hangover headache, has been recognized for at least nineteen-hundred years, yet the mechanism behind this headache trigger is poorly understood (Dueland, 2015; Leith, 2015; Panconesi, 2016). Migraineurs experience more severe headaches with much less alcohol than non-migraineurs, suggesting that while the physiology responsible for DEIH is common, underlying pathophysiology is uniquely present in migraine patients (Anon, 2004). Understanding these mechanisms may reveal pathological systems unique to amongst migraineurs and potentially targetable. We used two rat models of trigeminal sensitivity with migraine-like features to study this common form of headache.

It was previously shown that a rat model of trigeminal sensitivity is susceptible to ethanol-induced trigeminal sensitivity (Maxwell et al., 2010). Similar to DEIH in humans, these rats experience increased trigeminal sensitivity 4-6 hours following ethanol treatment (Anon, 2004). Ethanol is metabolized into acetaldehyde which is then quickly metabolized into acetate (Zimatkin et al., 2006; Jiang et al., 2013). Since ethanol and acetaldehyde are both absent during the onset of pain, it was suggested and later confirmed that acetate was the key ethanol metabolite responsible for DEIH in this rat model (Tsukamoto et al., 1989). This study also illustrated that caffeine, a non-specific adenosine receptor antagonist, could prevent ethanol-induced trigeminal sensitivity, suggesting that adenosine played a role in DEIH.

In the brain, acetate is preferentially utilized by astrocytes where its converted into acetyl-coenzyme A in an ATP-dependent reaction, resulting in a residual AMP molecule that’s readily converted into adenosine and released to act at purinergic receptors throughout the brain (Carmichael et al., 1991; Pascual et al., 2005; Panatier et al., 2011; Zorec et al., 2012; Jiang et al., 2013). Serum-adenosine levels increase after ethanol ingestion in humans (Nagy, 1992) and extracellular adenosine concentrations increase in rats during microdialysis perfusion of ethanol in the brain (Sharma et al., 2010). In fact, the purinergic system has been hypothesized to play a role in migraine and pain (Cieślak et al., 2015; Fried et al., 2017).

Similar to migraine headache, DEIH includes throbbing pain with phono- and photophobia that is exacerbated with physical movement (García-Azorín et al., 2020). The headache phase of migraine is thought to be caused by activation of dural C and Aδ nociceptors of the trigeminal ganglion that project to the trigeminal nucleus caudalis (TNC) in the brainstem (Burstein et al., 2011). Two rat models of chronic trigeminal sensitivity with migraine-like characteristics currently exist to study migraine pathophysiology. The recurrent inflammatory stimulation model (IS rats) is induced by infusing inflammatory agents onto the dura through an affixed cannula 3x/week for four weeks (Oshinsky and Gomonchareonsiri, 2007). These animals transition to a steady-state of chronic trigeminal sensitivity that outlasts the final infusion for months. The spontaneous trigeminal allodynia model (STA rats) features inheritable primary trigeminal allodynia (Oshinsky et al., 2012; Munro et al., 2018). Similar to migraineurs who experience a combination of headache-free and headache days with fluctuating intensity, these animals feature episodically fluctuating trigeminal sensitivity. The goal of using two different rodent models of headache, featuring different sources of trigeminal pain/sensitization, is to demonstrate that the role of adenosine in DEIH is independent of the particular features of these individual models

We investigated acetate modulation of the adenosinergic system in the IS and STA rat models to explore the mechanisms behind DEIH. To determine the adenosine receptor subtype responsible for this form of trigeminal sensitivity, we pharmacologically manipulated adenosine receptor signaling during acetate treatment while measuring trigeminal sensitivity behaviorally. We then used microdialysis and HPLC to measure adenosine, glutamate, and glutamine in response to perfusion of acetate in the TNC. Our data support the hypothesis that acetate derived from ethanol metabolism induces accumulation of extracellular adenosine which produces trigeminal sensitivity through modulation of glutamateric signaling via adenosine A_2A_ receptors within the TNC. This study is the first to identify physiological consequences of acetate on adenosinergic systems within the trigeminal pain system.

## MATERIALS AND METHODS

A combination of male Sprague Dawley rats from Charles River (used to induce the IS model) and an in-house colony originating from Charles River (STA rats) (250-300 g, n = 168) were housed individually in a temperature-controlled environment under a 12-hour light/dark cycle. Animals were allowed access to food and water ad libitum. All procedures performed on the animals were approved by the Thomas Jefferson University Institutional Animal Care and Use Committee. Efforts were made to minimize animal numbers and suffering.

### Induction of the Inflammatory Soup Rats

#### Cannula implantation

Surgical procedures for the inflammatory stimulation model were previously described (Oshinsky and Gomonchareonsiri, 2007). Briefly, two weeks of habituation and training were performed prior to putting the rats under isoflurane anesthesia (3% induction, 1.5% maintenance) mixed with compressed air. A 3 mm wide craniotomy was performed above the junction of the superior sagittal and transverse sinuses to expose the dura. A plastic cap and stainless steel cannula (26 gauge, Plastics One Inc., Roanoke, VA, USA) was secured to the skull with a combination of small screws and dental cement. This was then sealed with an obdurator that was custom cut to extend just beyond the internal end of the cannula above the dura to prevent scar tissue formation that could obstruct the flow of compounds onto the dura. Rats recovered from surgery for one week, during which trigeminal pressure thresholds and weight were monitored to ensure the return to baseline levels.

#### Inflammatory stimuluation

Inflammatory stimulation of the dura with 0.1 mM prostaglandin E2 in 0.9% sterile saline (Sigma Aldrich, St Louis, MO) was performed in their home cage. Polyethylene tubing (PE50) was connected to a 30 µl Hamilton syringe and the exposed cannula. 25 µl of the IS was steadily infused over 5 minutes at an approximate rate of 5 µl per minute. Infusions were performed 3x/wk for a total of 12 infusions. Rats that develop chronic trigeminal sensitivity (< 4 g thresholds) during this infusion period that outlasts the final infusion for at least 1 week were used in these studies.

### Spontaneous Trigeminal Allodynia Rats

The spontaneous trigeminal allodynia rats in this paper are an in-house colony originating from Charles River representing the 17^th^ and 18^th^ generation of an inbred line of rats (Oshinsky et al., 2012). These animals experience spontaneous trigeminal allodynia similar to how migraineurs experience headache and headache-free days without the need for surgical or experimental manipulation. STA rats do not experience sensitivity in any area outside of the trigeminal region. These rats are monitored each day to assess their morning periorbital thresholds. Experiments are conducted on days when rats have morning thresholds less than 4 g.

### Tactile Sensory Testing

Rats, tested during the day, were trained and acclimated to a plastic tube restraint (inner diameter 8 cm, length 25 cm) before and after cannula implantation. Rats entered uncoaxed into this atraumatic restrainer to prevent the rats from walking away during sensory testing.

Periorbital pressure thresholds were assessed by applying von Frey monofilaments (Stoelting Co., Wood Dale, IL, USA) to both the left and right sides of the face over the medial portion of the eye. These von Frey hairs are calibrated nylon monofilaments that generate a reproducible buckling stress. Each monofilament is identified by manufacturer-assigned force values (10, 8, 6, 4, 2, 1.4, 1, 0.6, 0.4, 0.16, 0.07, 0.04, 0.02, 0.008 grams). The higher the value on the monofilament, the stiffer and more difficult it is to bend. For each time point, the right and left threshold data are recorded individually. The von Frey stimuli were presented in sequential descending order to determine the threshold of response. Several behaviors presented by the rat were considered a positive response: vigorously stroking its face with the ipsilateral forepaw, quickly recoiling its head away from the stimulus, or vocalizing. A force value is considered positive if the rat responds in a positive way to 2 of 3 trials of the von Frey monofilament. After a positive response, a weaker stimulus was presented. The threshold is considered the lowest force value that produces a positive response (2 out of 3 trials). Results are presented as the proportional change in periorbital threshold from baseline levels. Rats that did not respond to the 10 g stimulus were assigned 10 g as their threshold for analysis.

### Drug Treatment

After rats were confirmed to have developed chronic trigeminal sensitivity for at least 1 week following the last infusion of IS, animals were treated with ethanol, acetate, adenosine, adenosine antagonists/agonists, α-Cyano-4-hydroxycinnamic acid, or combination thereof at effective concentrations found throughout the literature. Rats had free access to water and standard rodent chow throughout the duration of all experiments. Periorbital thresholds were assessed prior to each treatment to establish baseline sensitivity. Periorbital thresholds were assessed hourly for 8 hours following ethanol (300 mg/kg i.p. Sigma, St Louis, MO) administration as previously performed (Maxwell et al., 2010). Periorbital thresholds were assessed each hour for 5 hours following acetate (60 mg/kg i.p. Sigma, St Louis, MO) administration. Caffeine (50 mg/kg i.p. Sigma, St Louis, MO) was administered 1.5 hours after acetate treatment as previously performed (Maxwell et al., 2010).

Adenosine receptor subtype-specific antagonists were administered [3mg/kg DPCX p.o., adenosine A_1_ receptor antagonist, Tocris, Bristol, UK] [0.1 mg/kg istradeffyline p.o., adenosine A_2A_ receptor antagonist, Tocris, Bristol, UK] [12.5 mg/kg alloxazine p.o., adenosine A_2B_ receptor antagonist, Sigma, St Louis, MO] [1 mg/kg MRS-1523 p.o., adenosine A_2b_ receptor antagonist, Sigma, St Louis, MO] 1.5 hours after acetate treatment in individual experiments. α-Cyano-4-hydroxycinnamic acid o.g. (4-CIN) (100 mg/kg) was administered 30 minutes prior to acetate treatment to ensure this competitive inhibitor was at the site of action before acetate.

To determine the local effects of adenosine receptor signaling on trigeminal pain, compounds were directly injected into the cisterna magna, an opening of the subarachnoid space which allows for direct access to the region of the brainstem which contains the trigeminal nucleus caudalis (TNC). Intracisternal injection of capsaicin induces c-Fos activation primarily in the TNC, validating the use of these injections to target the region of the brainstem containing the TNC (Mitsikostas et al., 1998, 1999). 10 µl of 12.5 µM adenosine (Sigma, St Louis, MO) or aCSF was injected into the cisterna magna. To target centrally acting A_2A_ receptors within the TNC 10 µl of 300 µM CV-1808 [adenosine A_2A_ /A_2B_ receptor agonist, Tocris, Bristol, UK] was injected into the cisterna magna 30 minutes following administration of alloxazine (12.5 mg/kg p.o.). Dose and time-points for each compound were selected based on literature and pilot studies.

### Acetate perfusion and microdialysis sampling of the trigeminal nucleus caudalus

EiCOM CX-I-12-01 brain probes (EiCOM USA, San Diego, CA) with a 12 mm guide cannula length, 1 mm artificial cellulose membrane length, 50,000 Dalton cutoff, and a 0.22 mm outer diameter had their individual recovery rates determined prior to the experiment with adenosine, glutamate and glutamine standards (Sigma Aldrich, St Louis, MO). Recovery rates varied between 5-10% for adenosine. Probes were placed into the trigeminal nucleus caudalis (TNC) of anesthetized rats -2.6 to -2.9 mm from obex and 1.7 to 1.9 mm lateral to the midline. The probe’s inlet was connected with PE10 tubing to a CMA 110 Liquid Switch and then to a 2.5 mL glass syringe mounted on a CMA/100 microinjection pump. The probe’s outlet was connected to a CMA 170 refrigerated fraction collector (CMA Microdialysis AB, North Chelmsford, MA, USA) with PE10 tubing. The dialysis system was perfused at 0.7 μL/min with sterile, pyrogen-free artificial extracellular spinal fluid (aCSF; composition in mM/L: 135 NaCl; 3 KCl; 1 MgCl_2_; CaCl_2_; pH = 7.2). Probes were inserted into anesthetized animals 2.5 hours prior to experiment to allow neurotransmitters to settle to baseline levels. Baseline levels of adenosine, glutamate, and glutamine were confirmed to occur after 2 hours following probe insertion with no statistical change in neurotransmitter levels for the next 6 hours (n = 4). Samples were collected continuously every 20 minutes and collection began immediately after probe insertion.

Experimental concentrations were calibrated for each probe by using the recovery rate to calculate the acetate concentration needed to obtain an extracellular concentration of 2 mM, 10 mM, 20 mM, or 40 mM acetate in the TNC during perfusion (i.e., if 5% recovery rate, then perfusate contained 40 mM of acetate to achieve 2 mM of acetate in the TNC). The average recovery rate across all probes used was 6.1 ± 0.4%. Syringes of aCSF containing these calculated levels of acetate (Sigma-Aldrich, St Louis, MO) were mounted onto the microinjection pump. Three samples (i.e. 60 minutes of collection) were obtained in a sequentially step-wise increasing manner for each concentration by switching the inlet lines with the CMA 110 Liquid Switch, accounting for dead space of tubing.

Similar calculations using the recovery rate for each probe were used for experiments containing 4-CIN (Sigma-Aldrich, St Louis, MO) and istradeffyline (Tocris, Bristol, UK) within the perfusate. For these experiments, aCSF containing sufficient concentrations to achieve 200 µM 4-CIN or 5 nM of istradeffyline was perfused for 40 minutes prior to acetate exposure. These concentrations of 4-CIN and istradeffyline were sustained throughout the entire experiment as they were included with acetate concentrations necessary to obtain 10 mM and 40 mM local acetate concentrations in the TNC.

At the end of the experiment, animals were administered 0.5 mL of Euthasol via i.p. injection. The probe was removed and stored in distilled water. The brain and spinal cord were removed and stored in 4% paraformaldehyde for later analysis to check the position of the probe via cryosectioning.

### HPLC Measurement of Amino Acids and Adenosine

The amino acid content (glutamate and glutamine) of each sample was analyzed via high-performance liquid chromatography (HPLC) using a binary gradient and pre-column derivatization of O-phthal aldehyde (OPA) with fluorescence detection.

Samples were diluted (4 μL aCSF + 6 μL dialysate) and a 1:2 sample to reagent ratio was used (10 μL sample + 20 μL OPA). After a 60 second reaction, 20 μL of the sample-OPA mixture was auto-injected into an Agilent Zorbax Eclipse AAA column (150 x 4.6 mm; 5 μm particle size). A binary gradient of mobile phase A (40 mM sodium phosphate monobasic; pH = 7.4) and mobile phase B (45% acetonitrile; 45% methanol; 10% water) with a flow rate of 1.5 mL/min was used for separation.

Adenosine content of each sample was separated via HPLC and analyzed with a UV detector using previously developed methods (Sharma et al., 2010). 10 μl of dialysate sample was injected into the HPLC system containing a mobile phase of 8 mM NaH_2_PO_4_ and 8% methanol (pH = 4) at a flow rate of 80 μl/min. Adenosine was separated with a microbore column (1 × 100 mm; MF-8949; BASi, West Lafayette, IN) and detected with a UV detector (Model SPD-10Avp/10AVvp, Shimadzu Scientific Instruments, Columbia MD) at a 258 nm wavelength.

Column temperatures were maintained at 30°C. EZChrom Elite version 3.1.6 software was used to determine concentrations of extracellular neurotransmitters by comparing retention time and area under the peak to known amounts of standards

### Experimental Design and Statistical Analysis

All statistical analyses were performed using SPSS version 19. Sufficient power was confirmed with a power analysis for these studies using PASS software (NCSS, Kaysville, UT) to limit the number of animals needed for these experiments. Periorbital thresholds at individual time points were analyzed using a one-way ANOVA to determine drug effects on trigeminal sensitivity. One-way repeated measure ANOVA was used to compare the effect of acetate on neurotransmitter release and a one-way ANOVA was used at each acetate concentration to compare groups. Bonferroni post hoc tests were performed where applicable. A t-test was used to compare basal levels of neurochemicals.

## Results

We used two rat models of trigeminal sensitivity that feature several migraine-like traits: the recurrent inflammatory stimulation (IS) rat model and the spontaneous trigeminal allodynia (STA) rat model. Both systems model the trigeminal sensitivity (IS-chronic sensitivity, STA-episodic sensitivity), phonophobia, sensitivity to migraine triggers, and similar efficacious responses to migraine treatments (Oshinsky and Luo, 2006; Maxwell et al., 2010; Fried et al., 2014; Oshinsky et al., 2014; Munro et al., 2018). Trigeminal sensitivity was measured in male STA (n = 54) and IS rats (n = 107) in response to acetate treatment (60 mg/kg i.p.) while pharmacologically manipulating adenosine receptor signaling to determine the adenosine receptor subtype that is responsible for acetate-induced trigeminal sensitivity in the delayed ethanol-induced headache (DEIH). Extracellular adenosine, glutamate, and glutamine were measured with microdialysis and HPLC in the trigeminal nucleus caudalis (TNC) in response to local acetate perfusion to assess the mechanisms behind acetate-induced trigeminal sensitivity in DEIH.

### Acetate induces sensitivity in two rat models of trigeminal sensitivity

Ethanol treatment (300 mg/kg i.p.) of IS rats caused a biphasic effect on trigeminal pain thresholds (Fig. 1A). Initially, an analgesic effect was produced 1-2 hr post ethanol treatment as expressed by significantly higher periorbital thresholds in comparison to saline treated animals at the 1 hr (p = 0.011) and 2 hr (p = 0.031) time points. Animals then experienced an increase in trigeminal sensitivity as expressed by significantly lower periorbital thresholds during the 4 hr (p = 0.013) and 5 hr (p = 0.008) time points in comparison to saline treated animals (Fig. 1A). Ethanol is metabolized into acetaldehyde by alcohol dehydrogenase which is subsequently metabolized to acetate by aldehyde dehydrogenase. Previous pharmacological studies illustrated that acetate is the key pro-nociceptive metabolite of ethanol. Bypassing unrelated effects of ethanol by directly treating IS rats with acetate (60 mg/kg i.p.) increased trigeminal sensitivity as expressed by significantly lower periorbital thresholds during the 1 hr (p = 0.032), 2 hr (p < 0.001), 3 hr (p < 0.001) and 4 hr (p = 0.014) hour time points in comparison to saline treated animals (Fig. 1B). These treatment concentrations (60 mg/kg i.p. acetate and 300mg/kg i.p. ethanol) result in similar serum-acetate levels in the rat (approximately 1 mM) (Maxwell et al., 2010).

**Figure 1:**
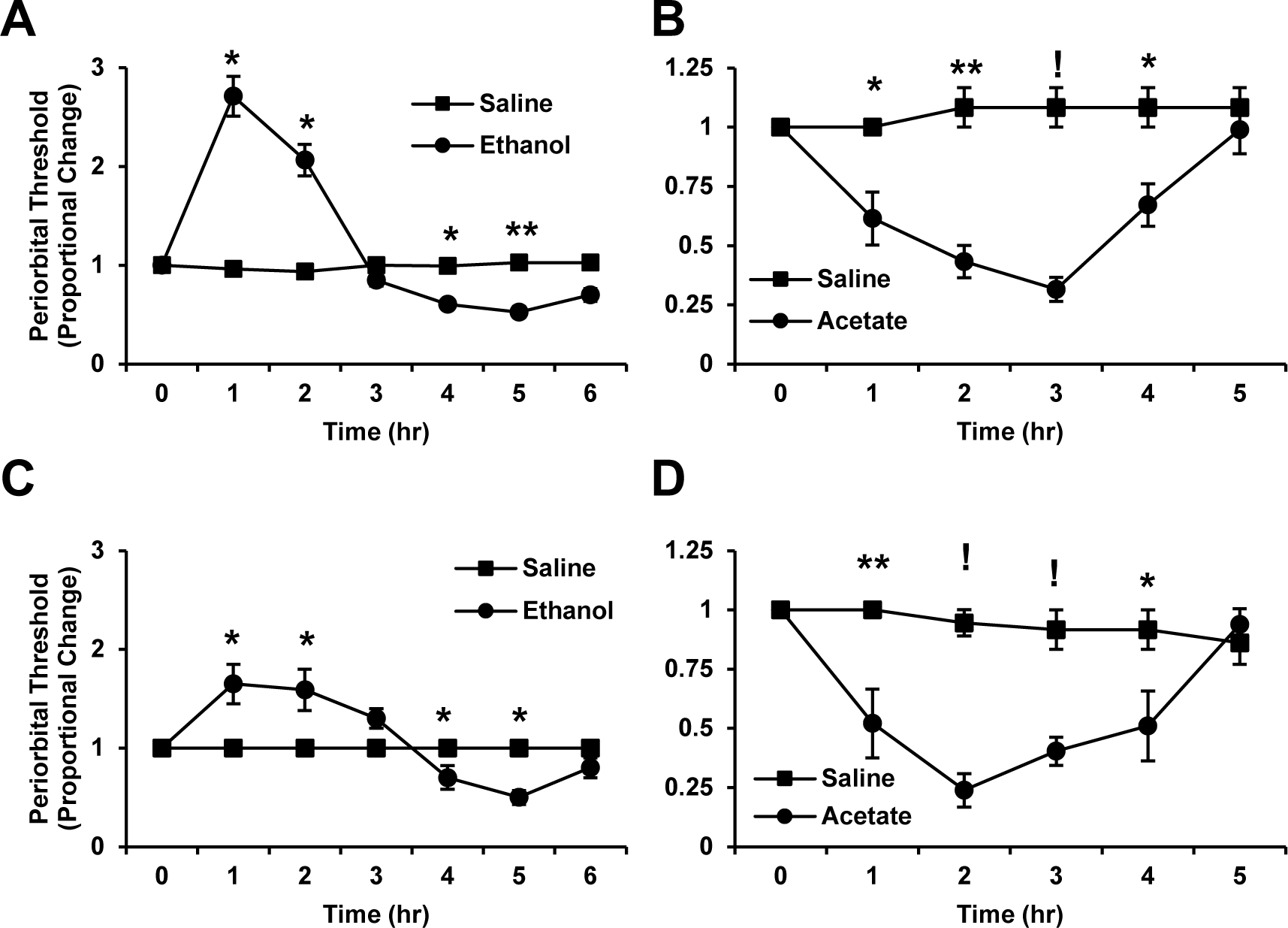
Ethanol and acetate treatment of IS and STA rat models. Comparison of the effect of ethanol or acetate vs saline on periorbital von Frey hair withdrawal thresholds in the IS and STA rat models of trigeminal pain as measured by proportional change in threshold from baseline. A) Ethanol treatment (300 mg/kg i.p., n = 9) vs saline (n = 9) in IS rats. B) Acetate treatment (60 mg/kg i.p., n = 7) vs saline (n = 5) in IS rats. C) Ethanol treatment (300 mg/kg i.p., n = 10) vs saline (n = 10) in STA rats. D) Acetate treatment (60 mg/kg i.p., n = 7) vs saline (n = 6) in STA rats. (*, *p* < 0.05) (**, p < 0.01) (!, p < 0.001).

We confirmed that ethanol and acetate also cause increased trigeminal sensitivity in the STA rat model of trigeminal pain. In this model, ethanol (300 mg/kg i.p.) also produced a biphasic effect on trigeminal sensitivity. An analgesic effect initially occurred as seen by significantly higher periorbital thresholds at the 1 hr (p = 0.043) and 2 hr (p = 0.045) time points in comparison to the saline treated animals. This was followed by increased trigeminal sensitivity as expressed by significantly lower periorbital thresholds at the 4 hr (p = 0.042) and 5 hr (p = 0.039) time points in comparison to saline treated animals (Fig. 1C). Acetate treatment (60 mg/kg i.p.) caused trigeminal sensitivity as seen by significantly lower periorbital thresholds during the 1 hr (p = 0.007), 2 hr (p < 0.001), 3 hr (p < 0.001), and 4 hr (p = 0.034) time points in comparison to saline treated animals (Fig. 1D).

### Competitive inhibition of monocarboxylate transport prevents acetate-induced trigeminal sensitivity

Acetate is primarily utilized as an energy substrate by astrocytes within the brain since neurons do not import it (Waniewski and Martin, 1998; Wyss et al., 2011). To determine whether acetate entry into astrocytes is required for acetate’s effect on trigeminal sensitivity, STA and IS rats were treated with alpha-cyano-4-hydroxycinnamate (4-CIN), a monocarboxylate transporter (MCT) competitive inhibitor, thirty minutes prior to acetate treatment. Treatment of IS rats with 4-CIN (100 mg/kg p.o.) attenuated the acetate-induced trigeminal sensitivity as seen by a significantly higher threshold in the 4-CIN co-treated animals in comparison to the acetate only animals at the 1 hr (p = 0.003), 2 hr (p < 0.001), 3 hr (p < 0.001), and 4 hr (p = 0.03) time points (Fig. 2A).

**Figure 2:**
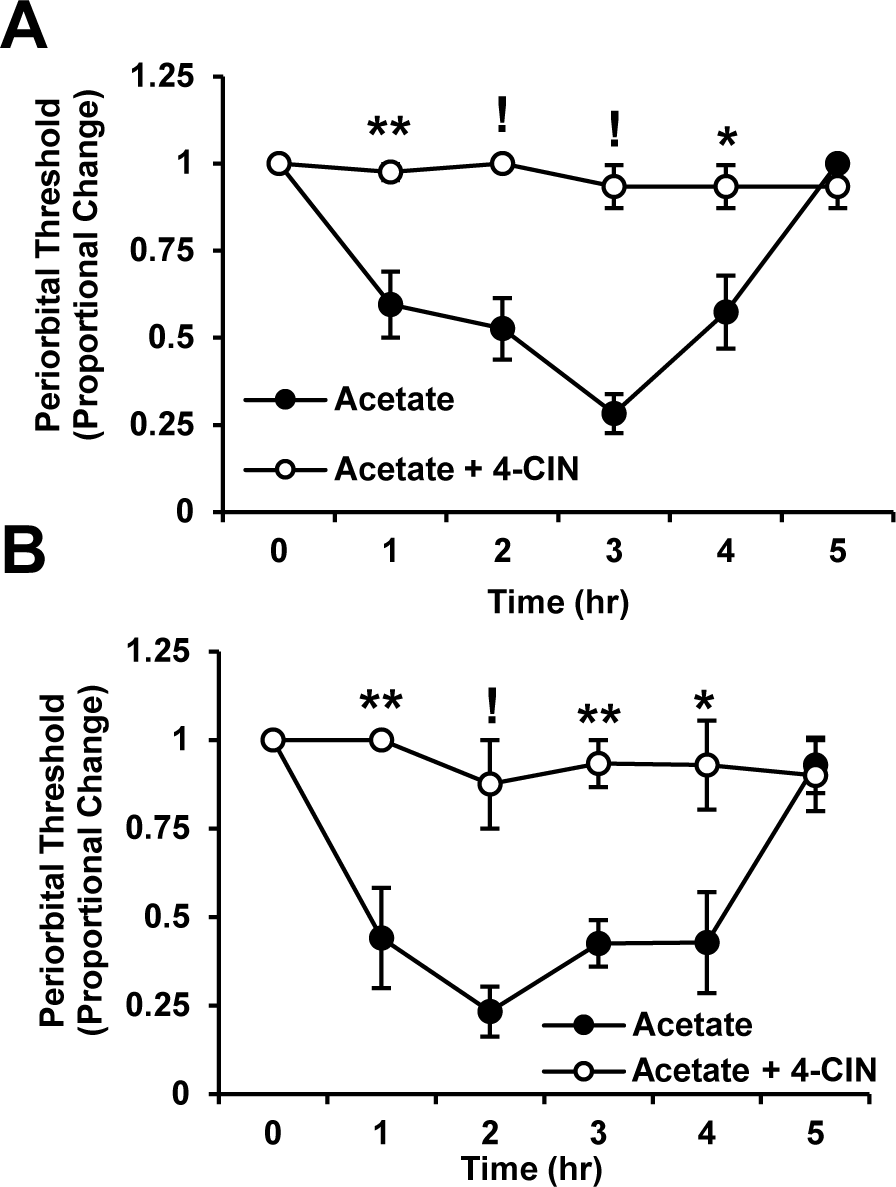
4-CIN treatment of acetate-treated IS and STA rats. Measurement of the effect of α-Cyano-4-hydroxycinnamic acid (4-CIN), a competitive MCT inhibitor, on acetate-induced trigeminal sensitivity as measured by proportional change in periorbital von Frey hair withdrawal thresholds from baseline. IS rats (A) and STA rats (B) were administered 4-CIN (100 mg/kg p.o.) 30 minutes prior to acetate treatment (60 mg/kg i.p.) and compared to the acetate-only group. Each group consists of 6 animals. (*, *p* < 0.05) (**, p < 0.01) (!, p < 0.001).

Treatment with 4-CIN (100 mg/kg p.o.) also attenuated the acetate-induced trigeminal sensitivity in STA rats as seen by a significantly higher threshold in the 4-CIN co-treated animals in comparison to the acetate only animals at the 1 hr (p = 0.003), 2 hr (p < 0.001), 3 hr (p < 0.001), and 4 hr (p = 0.03) time points (Fig 2B). 4-CIN administered to IS or STA rats in the absence of acetate had no effect on thresholds across 5 hours (Sup. Fig. 1). These data suggest that acetate-induced trigeminal sensitivity requires acetate entry, and possibly metabolism, in astrocytes.

### Adenosine A_2A_ receptor activation is necessary for acetate-induced trigeminal sensitivity

Caffeine treatment, a non-specific adenosine receptor antagonist, prevents ethanol-induced trigeminal sensitivity in IS rats (Maxwell et al., 2010). To confirm adenosine receptor involvement in acetate-induced trigeminal sensitivity, IS rats were treated with caffeine (50 mg/kg i.p.) 1.5 hour following acetate treatment. This timing was used due to caffeine’s relatively short half-life in rats (1.2 hr) to ensure sufficient antagonism during the peak of acetate-induced trigeminal sensitivity (Nehlig, 1999). This also allowed us to confirm acetate’s initial effect on trigeminal thresholds during the 1 hr time point prior to drug treatment. Similar to caffeine’s effects on ethanol-induced trigeminal sensitivity, caffeine treatment attenuated acetate-induced trigeminal sensitivity as seen by a significantly higher threshold in the caffeine co-treated animals in comparison to the acetate only animals at the 2 hr (p = 0.009), 3 hr (p < 0.001), and 4 hr (p = 0.006) time points (Fig. 3A). Animals receiving caffeine co-treatment had similar thresholds to acetate-only treated animals at the 1 hr time point, prior to caffeine treatment (p = 0.654). These data suggest that adenosine signaling plays a role in acetate-induced trigeminal sensitivity.

**Figure 3:**
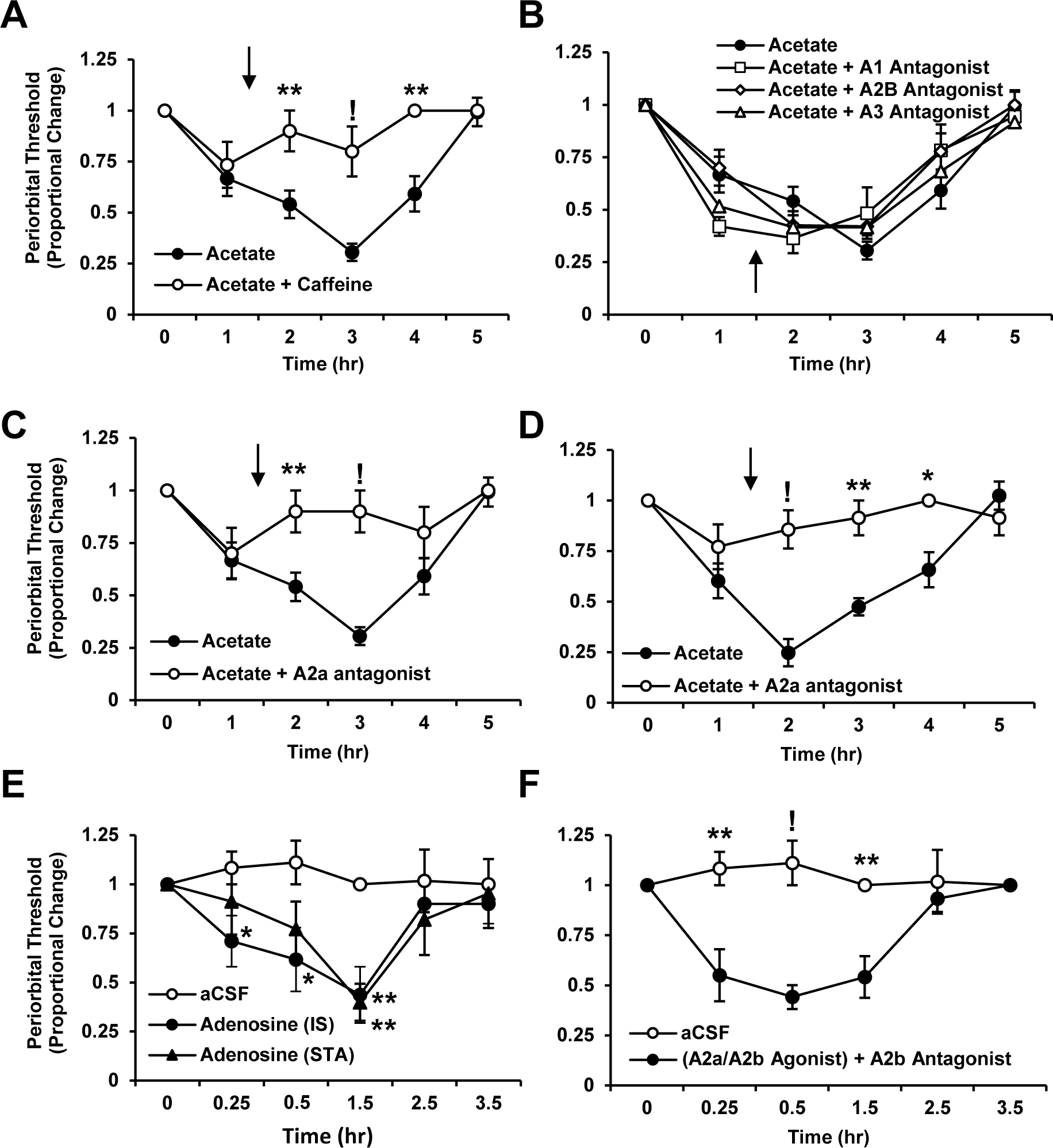
Adenosine receptor subtype involvement in acetate-induced trigeminal sensitivity in the IS and STA rats. Measurement of the effect of adenosine receptor modulation on acetate-induced trigeminal sensitivity as measured by proportional change in periorbital von Frey hair withdrawal thresholds from baseline in IS and STA rats. A) The effect of treatment with caffeine (50 mg/kg i.p.) 1.5 hours after acetate treatment (60 mg/kg i.p.) compared to acetate-only in IS rats. B) The effect of treatment with the adenosine A_1_ receptor antagonist (DPCX 3 mg/kg p.o.), the A_2B_ receptor antagonist (alloxazine 12.5 mg/kg p.o.), or the A_3_ receptor antagonist (MRS-1527 1 mg/kg p.o.) 1.5 hours after acetate treatment (60 mg/kg i.p.) compared to acetate-only in IS rats. C) The effect of treatment with istradefylline (0.1 mg/kg p.o.), an A_2A_ antagonist, 1.5 hours after acetate treatment (60 mg/kg i.p.) compared to acetate-only in IS rats. D) The effect of treatment with istradefylline (0.1 mg/kg p.o.), an A_2A_ antagonist, 1.5 hours after acetate treatment (60 mg/kg i.p.) compared to acetate-only in STA rats. E) The effect of intracisternal injection of adenosine (10 µl, 12.5 µM) compared to aCSF in IS and STA rats. F) The effect of intracisternal injection of an adenosine A_2A_/A_2B_ receptor agonist (CV-1808; 10 µl, 300 µM) 30 minutes following administration of the A_2B_ receptor antagonist (alloxazine 12.5 mg/kg p.o.) compared to aCSF in IS rats. Solid arrow in A, B, C, and D indicates the time when the antagonist is administered. To increase visual clarity for comparison of groups, the acetate-only groups in A, B, and C are the same data, but all groups were compared with a one-way repeated measures ANOVA at the same time. For the same reasons, the aCSF groups in E and F are the same, but all groups were compared with a single repeated measures ANOVA. Acetate groups consist of 10 animals. All other groups consist of 6 animals each. (*, *p* <0 .05) (**, p < 0.01) (!, p < 0.001).

To determine which adenosine receptor subtype (A_1_, A_2A_, A_2B_, and A_3_) is primarily responsible for acetate-induced trigeminal sensitivity, IS rats were treated 1.5 hours after acetate administration with adenosine receptor subtype-specific antagonists and also compared to the animals treated only with acetate. For easier visualization of these comparisons, we separated this experiment into two figure panels (Fig. 3B and Fig. 3C) and thus the acetate group is the same as that seen in Fig. 3A, but all groups in Fig. 3A, 3B, and 3C were compared with a single one-way repeated measures ANOVA. Co-treatment with the adenosine A_1_ receptor antagonist (DPCX 3 mg/kg p.o.), the A_2B_ receptor antagonist (alloxazine 12.5 mg/kg p.o.), or the A_3_ receptor antagonist (MRS-1527 1 mg/kg p.o.) did not attenuate the acetate-induced trigeminal sensitivity in IS rats as seen by no significant differences in periorbital thresholds between the antagonist treated groups and the acetate-only group (Fig. 3B) (Jacobson and Gao, 2006).

Treatment with an A_2A_ receptor antagonist (istradefylline 0.1 mg/kg p.o.), however, attenuated acetate-induced trigeminal sensitivity as seen by a significantly higher threshold in the istradefylline co-treated animals in comparison to the acetate only animals at the 2 hr (p = 0.009) and 3 hr (p < 0.001) time points in the IS rats (Fig. 3C). Animals receiving istradefylline co-treatment had similar thresholds to acetate-only treated animals at the 1 hr time point, prior to istradefylline treatment (p = 0.826). We confirmed similar effects of istradefylline co-treatment in the STA rats where the istradefylline co-treated animals had significantly higher thresholds than then acetate-only group at the 2 hr (p < 0.001), 3hr (p = 0.002), and 4 hr (p = 0.03) time points (Fig. 3D). Istradefylline administered to IS or STA rats in the absence of acetate had no effect on thresholds across 5 hours (Sup. Fig. 1). These data suggest that acetate is inducing trigeminal sensitivity via adenosine A_2A_ receptors.

Following ethanol ingestion, acetate is produced throughout the entire body (Tsukamoto et al., 1989). Thus, acetate’s impact on trigeminal pain via adenosine receptor signaling may occur peripherally at primary afferents or centrally within the TNC, containing second order neurons. To determine if central adenosine receptor signaling is sufficient for trigeminal pain, we directly injected adenosine into the cisterna magna, an opening of the subarachnoid space, which allows for noninvasive access to the region of the brainstem containing the TNC. Intercisternal injections have previously been used to specifically target the TNC (Mitsikostas et al., 1998, 1999). For easier visualization of these comparisons, we separated this experiment into two figure panels (Fig. 3E and Fig. 3F) and thus the aCSF group is the same in both panels, but all groups in Fig. 3E and 3F were compared with a single one-way repeated measures ANOVA.

Intracisternal injection of adenosine (10 µl, 12.5 µM) in IS rats induced trigeminal sensitivity as seen by significantly lower thresholds in the adenosine treated animals in comparison to aCSF treated animals at the 0.25 hr (p = 0.034), 0.5 hr (p = 0.029), and 1.5 hr (p = 0.002) time points (Fig. 3E). This was confirmed in STA rats where significantly lower thresholds were seen in the adenosine treated animals in comparison to aCSF treated animals at the 1.5 hr (p < 0.001) time point (Fig. 3E).

To confirm the role of the A_2A_ receptors within the TNC, an adenosine A_2A_ /A_2B_ receptor agonist (CV-1808; 10 µl, 300 µM) was directly injected into the cisterna magna of IS rats thirty minutes following administration of the A_2B_ receptor antagonist (alloxazine 12.5 mg/kg p.o.) since no reliable A_2A_ exists. This allowed for selective stimulation of centrally-acting A_2A_ receptors within the TNC. This selective targeting of A_2A_ receptors induced trigeminal sensitivity as seen by significantly lower thresholds in the “A_2A_-agonist” animal group in comparison to aCSF treated animals at the 0.25 hr (p = 0.006), 0.5 hr (p < 0.001), and 1.5 hr (p = 0.001) time points (Fig. 3F).

### Acetate perfusion in the TNC induces dose-dependent increase in extracellular adenosine, glutamate, and glutamine concentrations

Microdialysis and HPLC was used to quantify the effect of central acetate perfusion on extracellular adenosine concentrations within the TNC of naive and IS rats. Local perfusion of acetate into the TNC with the use of microdiaylsis as opposed to systemic i.p. administration allowed for acetate exposure directly to the local brain parenchyma of the TNC. Use of acetate instead of ethanol allowed us to bypass the complicating factors of ethanol effects unrelated to pain. Calculating the *in vitro* recovery of each microdialysis probe, acetate concentrations were estimated in the perfusate for each experiment to obtain 0 mM, 2 mM, 10 mM, 20 mM, or 40 mM acetate around the vicinity of the probe tip in the TNC. After obtaining a stable baseline for 2-3 hours following probe insertion (this baseline is used to calculate fold change in each neurochemical), each concentration of acetate was sequentially perfused for 1 hour at a flow rate of 0.7 µl/min with 14 µl of microdialystate collected every 20 minutes (i.e., following baseline, 2 mM acetate is perfused for 1 hr, followed by 10 mM for 1 hr, followed by 20 mM for 1 hr, followed by 40 mM for one hr). 10 µl of each microdialysate was separated with HPLC and analyzed with a UV detector to measure extracellular adenosine concentrations (Sharma et al., 2010). The remaining 4 µl was separated with HPLC and used to determine extracellular concentrations of glutamate and glutamine via OPA-conjugated fluorescence detection. Fold change is calculated as the average fold change for each acetate concentration (i.e., the reported fold change is the average of the three samples collected during each acetate concentration).

Acetate perfusion induced a dose-dependent increase of extracellular adenosine in both IS and naive rats. The IS rats, however, experienced significantly larger increases of extracellular adenosine with perfusion of 2 mM (1.5 ± 0.2 fold in IS rats vs 0.9 ± 0.1 fold in naive rats, p = 0.035), 10 mM (2.7 ± 0.5 fold in IS rats vs 1.4 ± 0.1 fold in naive rats, p = 0.041), and 40 mM acetate (8.1 ± 1.8 fold in IS rats vs 2.5 ± 0.6 fold in naive rats, p = 0.025) (Fig. 4A). Perfusion of 20 mM acetate increased extracellular adenosine to similar levels in the IS rats (4.7 ± 1.1 fold) and naive rats (2.3 ± 0.5 fold) (Fig. 4A). Using each microdialysis probe’s individual recovery rate, estimation of basal adenosine concentrations within the TNC were calculated and were not statistically different between IS rats (159 ± 31 nM) and naive rats (188 ± 54 nM) (Fig. 4B).

**Figure 4:**
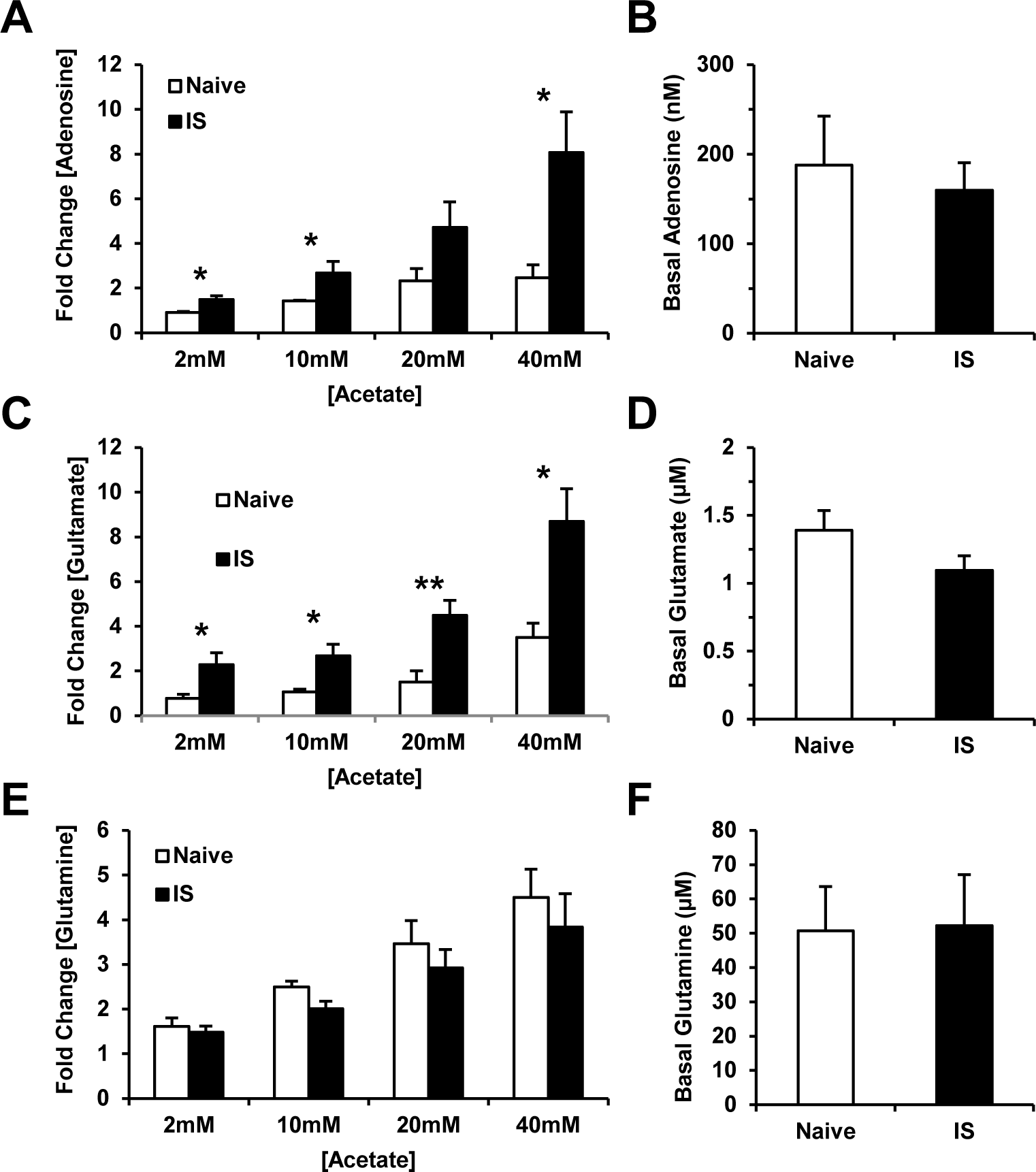
Measurement of extracellular adenosine, glutamate, and glutamine in response to acetate perfusion in the TNC. Microdialysis and HPLC analysis of acetate perfusion within the trigeminal nucleus caudalis (TNC) of IS rats induces greater increases in extracellular adenosine and glutamate, but not glutamine, in comparison to naive rats. A) Average fold change from baseline in extracellular adenosine levels in the TNC of IS and naive rats in response to 2 mM, 10 mM, 20 mM, and 40 mM acetate perfusion. B) Calculated basal levels of extracellular adenosine in the TNC of IS and naive rats. C) Average fold change from baseline in extracellular glutamate levels in the TNC of IS and naive rats in response to 2 mM, 10 mM, 20 mM, and 40 mM acetate perfusion. D) Estimated basal levels of extracellular glutamate in the TNC of IS and naive rats. E) Average fold change from baseline in extracellular glutamine levels in the TNC of IS and naive rats in response to 2 mM, 10 mM, 20 mM, and 40 mM acetate perfusion. F) Estimated basal levels of extracellular glutamine in the TNC of IS and naive rats. Each group consists of 7 animals. (*, *p* < 0.05) (**, p < 0.01).

Extracellular glutamate within the dorsal horn or the TNC has been used as a neurochemical marker for pain, thus, we quantified changes in glutamate levels in response to acetate perfusion (Tao et al., 2005; Oshinsky and Luo, 2006; Oshinsky et al., 2014). A dose-dependent increase of extracellular glutamate in response to acetate concentration was observed in the both IS and naive rats. IS rats, however, experienced a significantly higher level of extracellular glutamate with perfusion of 2 mM (2.2 ± 0.5 fold in IS rats vs 0.76 ± 0.2 fold in naive rats, p = 0.035), 10 mM (2.7 ± 0.5 fold in IS rats vs 1.1 ± 0.3 fold in naive rats, p = 0.035), 20 mM (4.5 ± 0.7 fold in IS rats vs 1.5 ± 0.4 fold in naive rats, p = 0.009), and 40 mM (8.7 ± 1.5 fold in IS rats vs 3.5 ± 0.8 fold in naive rats, p = 0.02) acetate (Fig. 4C). Estimated basal glutamate concentrations within the TNC were not statistically different between IS rats (1.1 ± 0.1 µM) and naive rats (1.4 ± 0.2 µM) (Fig 4D).

Upon acetate utilization as an energy substrate by astrocytes, acetyl Co-A is formed which enters the Krebs cycle, producing α-ketoglutarate that is used to produce glutamate. This glutamate is converted to glutamine and released from astrocytes for neuronal uptake as part of the glutamate-glutamine cycling between astrocytes and neurons (Jiang et al., 2013). Measurement of extracellular glutamine, therefore, can serve as a surrogate marker of acetate utilization. Glutamine levels were quantified in response to acetate perfusion in IS and naive rats. Although a dose-dependent increase of extracellular glutamate in response to acetate concentration was observed, no significant difference was found with perfusion of 2 mM (1.5 ± 0.1 fold in IS rats, 1.6 ± 0.4 fold in naive rats), 10 mM (2.0 ± 0.2 fold in IS rats, 2.5 ± 0.5 fold in naive rats), 20 mM (2.9 ± 0.4 fold in IS rats, 3.5 ± 0.3 fold in naive rats), or 40 mM (3.8 ± 0.7 fold in IS rats, 4.5 ± 0.8 fold in naive rats) acetate between IS and naive rats (Fig. 4E). This suggested that both groups metabolize similar levels of acetate within the TNC during microdialysis perfusion. Estimated basal glutamine concentrations within the TNC were not statistically different between IS rats (5.2 ± 1.5 µM) and naive rats (5.1 ± 1.3 µM) (Fig. 4F).

### 4-CIN co-perfusion prevents the increase in adenosine, glutamate, and glutamine during acetate perfusion

To determine whether acetate entry into astrocytes is required for the production of adenosine, 4-CIN (200 µM) was included in the acetate-containing perfusate during microdialysis of IS rats. To allow 4-CIN to sufficiently block MCT transport, 4-CIN containing aCSF was perfused 20 minutes prior to exposure of perfusate containing both acetate (10 mM or 40 mM) and 4-CIN (200 µM). The presence of 4-CIN prevented the acetate-induced increase in extracellular adenosine concentrations as seen by a significantly lower fold change during perfusion of 10 mM (1.0 ± 0.1 fold with 4-CIN vs. 2.7 ± 0.5 fold without 4-CIN, p = 0.042) but not 40 mM (3.9 ± 0.1 fold with 4-CIN, 8.1 ± 1.8 fold without 4-CIN) acetate (Fig. 5A). A similar effect was seen in extracellular glutamate and glutamine. The presence of 4-CIN prevented the acetate-induced increase in extracellular glutamate concentrations as seen by a significantly lower fold change during perfusion of 10 mM (1.2 ± 0.2 fold with 4-CIN vs. 2.9 ± 0.4 fold without 4-CIN, p = 0.001) but not 40 mM (6.6 ± 1.7 fold with 4-CIN, 9.9 ± 1.1 fold without 4-CIN) acetate (Fig. 5C). The presence of 4-CIN prevented the acetate-induced increase in extracellular glutamine concentrations as seen by a significantly lower fold change during perfusion of 10 mM (1.0 ± 0.1 fold with 4-CIN vs. 2.7 ± 0.5 fold without 4-CIN, p = 0.009) but not 40 mM (2.0 ± 0.2 fold with 4-CIN, 3.8 ± 0.7 fold without 4-CIN) acetate (Fig. 5E).

**Figure 5:**
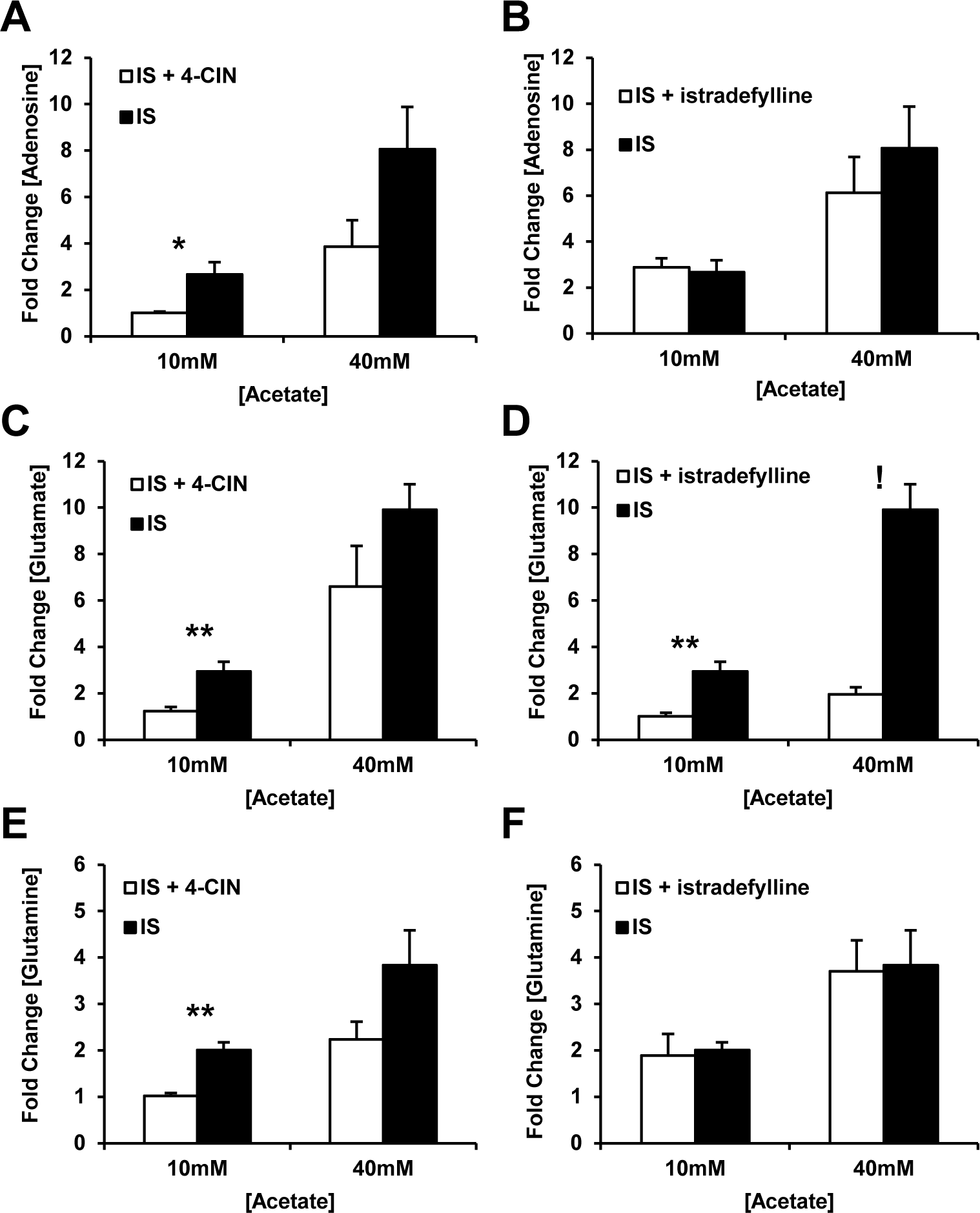
Measurement of acetate-induced changes in extracellular adenosine, glutamate, and glutamine in response to istradefylline or 4-CIN perfusion in the TNC. Microdialysis and HPLC analysis of acetate perfusion within the trigeminal nucleus caudalis (TNC) on extracellular concentrations of adenosine, glutamate, and glutamine of IS rats is modulated by istradefylline and 4-CIN. A) Average fold change from baseline in extracellular adenosine levels in the TNC of IS rats in response to 2 mM, 10 mM, 20 mM, and 40 mM acetate and 4-CIN (200 µM) perfusion. B) Average fold change from baseline in extracellular adenosine levels in the TNC of IS rats in response to 2 mM, 10 mM, 20 mM, and 40 mM acetate and istradefylline (5 nM) perfusion. C) Average fold change from baseline in extracellular glutamate levels in the TNC of IS rats in response to 2 mM, 10 mM, 20 mM, and 40 mM acetate and 4-CIN (200 µM) perfusion. D) Average fold change from baseline in extracellular glutamate levels in the TNC of IS rats in response to 2 mM, 10 mM, 20 mM, and 40 mM acetate and istradefylline (5 nM) perfusion. E) Average fold change from baseline in extracellular glutamine levels in the TNC of IS rats in response to 2 mM, 10 mM, 20 mM, and 40 mM acetate and 4-CIN (200 µM) perfusion. F) Average fold change from baseline in extracellular glutamine levels in the TNC of IS rats in response to 2 mM, 10 mM, 20 mM, and 40 mM acetate and istradefylline (5 nM) perfusion. Each group consists of 5 animals. (*, *p* < 0.05) (**, p < 0.01) (!, p < 0.001).

### Istradefylline co-perfusion prevents dose-dependent increase in glutamate but not adenosine or glutamine during acetate perfusion

To further determine the neurochemical mechanism behind acetate-induced trigeminal sensitivity and to determine if the increase in glutamate levels is a result of adenosine interaction with adenosine A_2A_ receptors, istradefylline (5 nM) was included in the perfusate during microdialysis in the TNC of IS rats. To allow sufficient antagonism of adenosine A_2A_ receptors, istradefylline was perfused through the microdialysis probe 20 minutes prior to exposure of perfusate containing both acetate and istradefylline (5 nM). Inclusion of istradefylline did not affect the acetate-induced increase in extracellular adenosine or glutamine during perfusion of 10 mM or 40 mM acetate, but it did significantly prevent acetate-induced increases in glutamate concentrations. Istradefylline presence did not significantly prevent the increase in extracellular adenosine concentrations during perfusion of 10 mM (2.8 ± 0.4 fold with istradefylline vs. 2.7 ± 0.5 fold without istradefylline) and 40 mM (6.1 ± 1.6 fold with istradefylline vs. 8.1 ± 1.8 fold without istradefylline) acetate (Fig. 5B). Istradefylline presence did significantly prevent the increase in extracellular glutamate concentrations during perfusion of 10 mM (1.0 ± 0.2 fold with istradefylline vs. 2.9 ± 0.4 fold without istradefylline, p = 0.005) and 40 mM (2.0 ± 0.3 fold with istradefylline vs. 9.9 ± 1.1 fold without istradefylline, p < 0.001) acetate (Fig. 5D). Inclusion of istradefylline did not significantly prevent the increase in extracellular glutamine concentrations during perfusion of 10mM (1.8 ± 0.5 fold with istradefylline vs. 2.0 ± 0.2 fold without istradefylline) and 40mM (3.7 ± 0.7 fold with istradefylline vs. 3.8 ± 0.7 fold without istradefylline) acetate (Fig. 5F).

## DISCUSSION

We identified adenosine as a critical component of the nociceptive effects of acetate in delayed ethanol-induced headache (DEIH). Adenosine A_2A_ receptor activation was essential for this effect in both rodent models, but the three other known adenosine receptor subtypes (A_1_, A_2B_, and A_3_) were not involved. This mechanism relies on MCT transport of acetate, likely into astrocytes which preferentially transport and metabolize acetate (Waniewski and Martin, 1998). Intracisternal injection of adenosine to target the TNC induced trigeminal sensitivity, suggesting that the mechanism behind DEIH is centrally-mediated (Panconesi, 2016). Acetate perfusion within the TNC induced a dose-dependent increase in adenosine, glutamate, and glutamine. In the IS rats, this effect was significantly greater for adenosine and glutamate, but not glutamine. This suggests that acetate utilization is similar in both groups, but that a physiological difference downstream of acetate conversion to acetyl-coA in the TNC of rats enhances the production of adenosine and glutamate. 4-CIN, an MCT competitive inhibitor, prevented this increase in adenosine, glutamate, and glutamine, supporting that acetate-induced release of these neurochemicals occur downstream of acetate transport. Istradefylline, an adenosine A_2A_ receptor antagonist, prevented the rise in glutamate but not adenosine or glutamine, suggesting that the increase in glutamate is downstream of adenosine A_2A_ receptor activation. Together, these data suggest a model for DEIH where adenosine derived from astrocytic acetate utilization activates adenosine A_2A_ receptors, which then modulates extracellular glutamate within the TNC to produce trigeminal sensitivity following ethanol exposure.

We find that adenosine A_2A_ receptor activation is responsible for modulation of extracellular glutamate levels in the TNC following systemic acetate treatment. Since glutamate is the principle excitatory neurotransmitter, its concentration is closely regulated via GLT1 (glutamate transporter 1) which is exclusively expressed on astrocytes to transport glutamate out of the synaptic cleft (Lehre and Danbolt, 1998; Anderson and Swanson, 2000). This is driven by a Na^+^ gradient maintained by the energetically-demanding Na^+^/K^+^ ATPase (NKA) on the cell surface (Zerangue and Kavanaugh, 1996; Rose et al., 2009). The reliance of this Na^+^ gradient and the process of maintaining it is so energetically demanding that co-compartmentalization of GLT-1, NKA, and mitochondria exists to allow rapid restoration of this critical Na^+^ gradient (Rose et al., 2009; Genda et al., 2011). Interestingly, modulation of this functional unit by inhibition of NKA activity, mitochondria, GLT1, or the Na+ concentration can affect glutamate transport (Keller et al., 1997; Li and Stys, 2001; Kawahara et al., 2002; Veldhuis et al., 2003; Köfalvi et al., 2020). Adenosine A_2A_ receptors physically associate with the α_2_ subunit of NKAs and their activation decreases NKA activity, modulating GLT1 function to decrease glutamate transport and increase its concentrations in the extracellular space levels (Matos et al., 2012, 2013). This mechanism may be responsible for the observation that extracellular glutamate concentrations can be modulated by the inhibition of adenosine A_2A_ receptors during acetate perfusion. An alternative mechanism may involve the excitatory effects of adenosine A_2A_ receptors on presynaptic neurons which can enhance synaptic transmission (Marchi et al., 2002).

Adenosine’s modulation of extracellular glutamate may play a key role in trigeminal pain. Extracellular glutamate within the TNC of IS rats has been used as a neurochemical marker of trigeminal pain (Oshinsky et al., 2014). Glutamate also plays a role in the spinal dorsal horn where modulation of glutamate is associated with other forms of chronic pain (Tao et al., 2005). Centrally-acting glutamate receptor antagonists are effective in reducing pain in animal models and humans while blocking excitatory amino acid transporters, increasing synaptic glutamate concentrations, produces pain (Hewitt, 2000; Liaw et al., 2005). Excess glutamate can lead to over-excitation of neuronal circuits within the dorsal horn or TNC, contributing to the development and maintenance of chronic pain states (Niederberger et al., 2003; Tao et al., 2005). Since adenosine A_2A_ receptor antagonism prevents the rise in glutamate and acetate-induced trigeminal sensitivity, adenosine likely modulates trigeminal pain via its effect on extracellular glutamate concentrations in the TNC during DEIH.

4-CIN, a MCT inhibitor, prevented acetate-induced trigeminal sensitivity in both models and prevented the increase in adenosine, glutamate, and glutamine in IS rats. This suggests that transport of acetate through MCTs is essential to the mechanisms behind acetate-induced trigeminal sensitivity in DEIH. Although monocarboxylic acid transport modulation can occur at the blood-brain barrier (BBB), we believe it is transport directly into astrocytes that is being modulated in this system (Muir et al., 1986; Terasaki et al., 1991; Waniewski and Martin, 1998; Hatazawa et al., 2010; Patel et al., 2010; Wyss et al., 2011; Jiang et al., 2013; DosSantos et al., 2014). Although MCTs play a vital role in transport of other monocarboxylic acids at the BBB, acetate is believed to not be regulated at the BBB, crossing by simple diffusion and is instead regulated at the cell surface of astrocytes (Terasaki et al., 1991; Waniewski and Martin, 1998; Patel et al., 2010). Thus, changes in BBB permeability seen in the IS model would not affect acetate transport, suggesting that 4-CIN regulation of acetate transport occurs at the astrocyte surface (Fried et al., 2018). 4-CIN does have affinity for other MCTs that modulate pyruvate or lactate transport in other cell types (Granja et al., 2013). We found, however, that 4-CIN alone does not change baseline trigeminal sensory thresholds, suggesting that its interaction with other MCT isoforms likely does not play a role in this process. 4-CIN’s ability to prevent acetate transport into astrocytes is further confirmed in that it prevented the formation of glutamine which is formed during acetate utilization.

We find that baseline adenosine concentrations within the TNC similar to concentrations observed in the rat hippocampus (120-200 nM), striatum (40-210 nM), and cortex (120 nM) (Latini and Pedata, 2001). Similar to human migraineurs who are more susceptible to DEIH than non-migraineurs, the acetate-induced adenosine rise is exacerbated in the IS rats in comparison to control animals. In fact, control animals need 20 mM of acetate perfused in the TNC to produce similar extracellular glutamate levels as IS rats experience with only 2 mM of acetate. This suggests that a physiological difference exists within the TNC of IS rats that may make them more susceptible to adenosine formation when exposed to acetate. Since there was no difference in glutamine production during acetate perfusion, this physiological difference is likely downstream of acetate conversion to acetyl-coA. This may be due to a decreased mitochondrial spare respiratory capacity that was identified in the mitochondria of brain slices from the TNC in both the IS and STA rat models (Fried et al., 2014; Fried and Oshinsky, 2015). Although this dysfunction does not impact basal levels of mitochondrial function, it may lower downstream processes involved in substrate utilization in active neuronal tissue which is already functioning at a level requiring 80% of the maximum capacity of mitochondrial output (Desler et al., 2012). Interestingly, mitochondrial dysfunction is associated with an increase in adenosine formation (Watanabe et al., 1983; Eltzschig et al., 2004; Duley et al., 2011). If this decrease in spare capacity within the TNC is present in astrocytes, the metabolic processes which control adenine nucleotide balance within the cell may be disrupted while those controlling acetate entry into the citric acid cycle (resulting in glutamine production) are not. Since DEIH can occur in non-migraineurs, it suggests that the mechanism of acetate-induced adenosine formation and subsequent modulation of extracellular glutamate is not unique to migraineurs or sensitized animals, but that it is exacerbated in these animals; potentially due to mitochondrial dysfunction or other physiological changes.

These results contribute to the migraine headache and alcohol fields at large. Adenosine plays a significant role in neurogenic inflammation, neuronal modulation, glial function, other forms of pain, changes in the blood-brain barrier and many other physiological processes that affect migraine (Guieu et al., 1998; Sawynok and Liu, 2003; Carman et al., 2011; Pedata et al., 2014). Notably, there is mounting evidence of adenosine as a critical player in the pathophysiology of migraine and headache (Fried et al., 2017). There is growing evidence suggesting that chronic pain and alcoholism may involve similar neurophysiological changes in overlapping brain regions involved in both disorders (Egli et al., 2012). Similar to DEIH, alcohol can produce hyperalgesia during the withdrawal period and conversely, chronic pain conditions are associated with alcohol misuse and dependence (Castillo et al., 2006; Gatch, 2009; Edwards et al., 2012). Adenosine is heavily involved in the development of alcoholism and may serve as the key neurophysiological modulator for the interplay between chronic pain and alcoholism (Ruby et al., 2010).

These studies expand on our previous work that identified acetate as the key metabolite of DEIH by providing further evidence that acetate’s effect on trigeminal pain is induced via adenosinergic mechanisms. We also provide evidence that the mechanism behind this effect likely involves modulation of extracellular glutamate levels via adenosine A_2A_ receptors. These data present, for the first time *in vivo*, evidence that supports adenosine A_2A_ receptor modulation of astrocytic glutamate transport (Genda et al., 2011; Matos et al., 2013). Additionally, these data illuminate adenosine as a potential key element to headache pathophysiology.

## AUTHOR CONTRIBUTIONS

NTF designed studies, acquired and analyzed data and the drafted manuscript. CRM designed and acquired data. JBH critically revised the manuscript. MBE made experimental, data interpretation and editorial contributions to the manuscript. MLO designed, analyzed, supervised the study and critically revised the manuscript. Each author gave final approval of the version to be published.

## Acknowledgments

The authors would like to thank Dr. Erin Seifert, Dr. Davide Trotti, and Dr. Stephen D. Silberstein for their guidance; Marnie Cooper, Leyla Murphy, Hugh Hekierski, Jr., Jessica Perino, Brittany Daiutolo, Ashley Tyburski, and the Jefferson Headache Center for their support. This work was supported by the National Institutes of Health (F31-AA017852 to CRM, R01-NS061571 to MLO, NIAAA K05-AA017261 to NTF), and the Migraine Research Foundation. The funding agencies had no role in study design, data collection and analysis, decision to publish, or preparation of the manuscript.

## Abbreviation

(IS): Inflammatory stimulation
(IS rats): inflammatory stimulation model
(STA rats): spontaneous trigeminal allodynia model
(DEIH): Delayed ethanol-induced headache
(TNC): Trigeminal nucleus caudalis

---

**Supplemental Figure 1:**
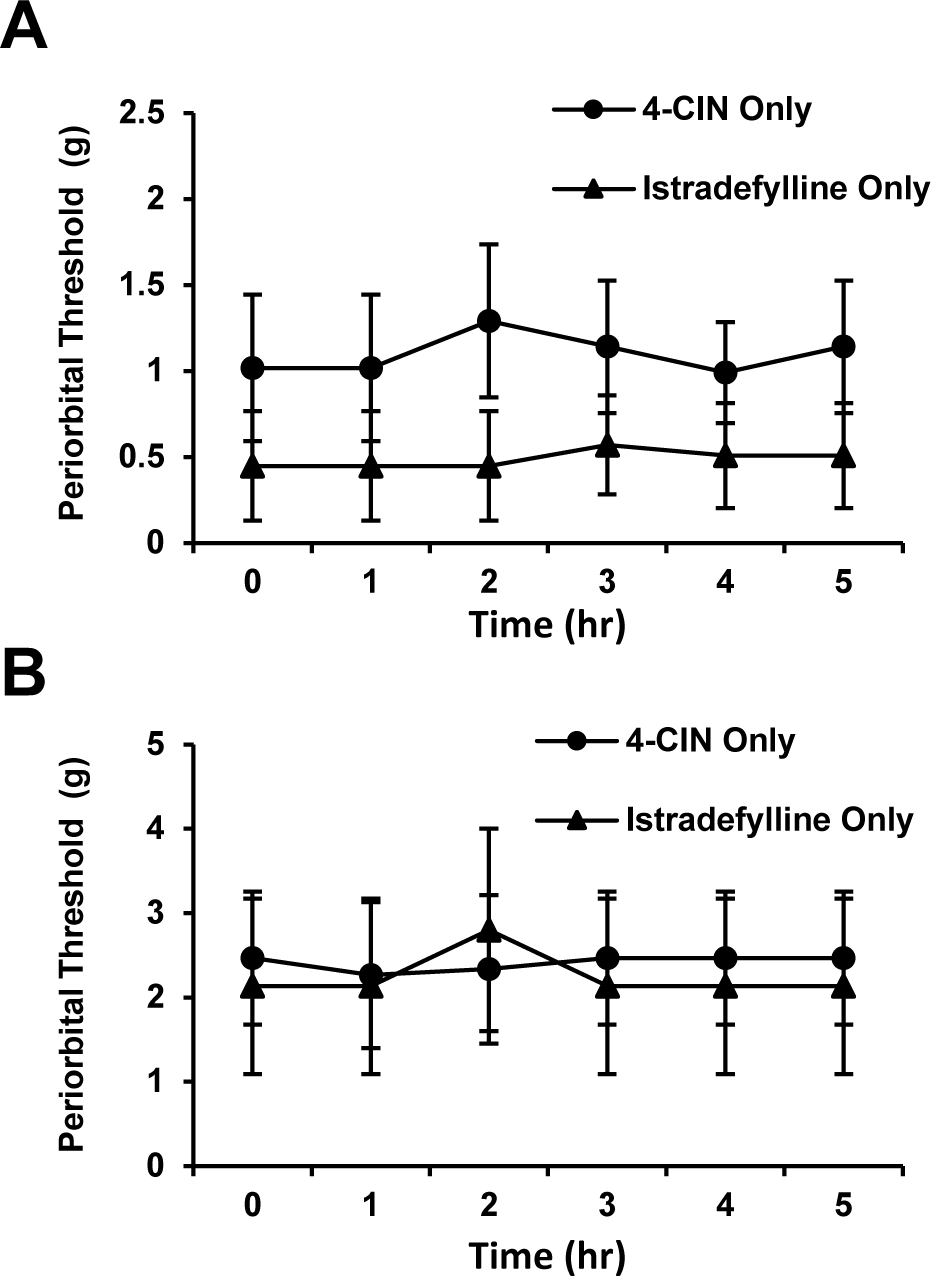
4-CIN and istradefylline alone treatments of IS and STA rats. Von Frey Hair periorbital thresholds (g) of IS and STA rats in response to 4-CIN and istradefylline treatment alone. IS rats (A) (n = 5) and STA rats (B) (n = 4) were administered either 4-CIN (100 mg/kg p.o.) or istradefylline (0.1 mg/kg p.o.) and periorbital thresholds were tracked over 5 hrs.

